# EEG microstate dynamics indicate a U-shaped path to propofol-induced loss of consciousness

**DOI:** 10.1101/2021.10.26.465841

**Authors:** Fiorenzo Artoni, Julien Maillard, Juliane Britz, Martin Seeber, Christopher Lysakowski, Lucie Bréchet, Martin R. Tramèr, Christoph M. Michel

## Abstract

It is commonly believed that the stream of consciousness is not continuous but parsed into transient brain states manifesting themselves as discrete spatiotemporal patterns of global neuronal activity. Electroencephalographical (EEG) microstates are proposed as the neurophysiological correlates of these transiently stable brain states that last for fractions of seconds. To further understand the link between EEG microstate dynamics and consciousness, we continuously recorded high-density EEG in 23 surgical patients from their awake state to unconsciousness, induced by step-wise increasing concentrations of the intravenous anesthetic propofol. Besides the conventional parameters of microstate dynamics, we introduce a new method that estimates the complexity of microstate sequences. The brain activity under the surgical anesthesia showed a decreased sequence complexity of the stereotypical microstates, which became sparser and longer-lasting. However, we observed an initial increase in microstates’ temporal dynamics and complexity with increasing depth of sedation leading to a distinctive “U-shape” that may be linked to the paradoxical excitation induced by moderate levels of propofol. Our results support the idea that the brain is in a metastable state under normal conditions, balancing between order and chaos in order to flexibly switch from one state to another. The temporal dynamics of EEG microstates indicate changes of this critical balance between stability and transition that lead to altered states of consciousness.

**Highlights:**

- EEG microstates capture discrete spatiotemporal patterns of global neuronal activity
- We studied their temporal dynamics in relation to different states of consciousness
- We introduce a new method to estimate the complexity of microstates sequences
- With moderate sedation complexity increases then decreases with full sedation
- Complexity of microstate sequences is sensitive to altered states of consciousness

## Introduction

Spontaneous mental activity is discontinuous and can be parsed into a series of conscious states manifesting themselves as discrete spatiotemporal patterns of global neuronal activity. Terms such as “pulses of consciousness” (James, 2007), “perceptual frames” (Efron, 1970), “neuronal workspace” (Baars, 1997; Dehaene et al., 1998), “heteroclinic channel” (Rabinovich et al., 2001) or “structure flow on manifolds” (Huys et al., 2014) describe the various concepts of parcellation of consciousness into sequential episodes - for reviews see (Deco et al., 2011; Meehan and Bressler, 2012; Michel and Koenig, 2018). For example, the neuronal workspace model suggests that discrete large-scale spatiotemporal neural activity patterns are transiently formed, remain for a certain amount of time, and then rapidly transition to a new co-activation pattern (Baars, 1997; Baars, 2002b; Dehaene et al., 2003). This model posits that only one global state exists at any moment in time and that conscious mentation emerges by the serial appearance of discrete states (Seth and Baars, 2005). A very similar chunking principle underlies the concept of “heteroclinic channels” (Rabinovich et al., 2001) that divide the mental activity into a chain of transient, metastable states. Metastability is a crucial principle that allows a system to spontaneously switch from one coordinated brain state to another, even in the absence of input (Deco and Jirsa, 2012; Jirsa et al., 1998; Tognoli and Kelso, 2014). Such flexible dynamics are important since conscious experiences are related to a rich and diverse repertoire of functional states which need to stabilize order and disorder, as unbalanced brain states can cause alterations in the global state of consciousness.

These brief periods of stable brain states switch from one to the other on the sub-second scale. Many behavioural studies have shown that the duration needed for conscious experience is in the range of 100 ms (Dehaene et al., 2003; Efron, 1970; Libet, 1981). On a neuronal level, similar durations have been measured for synchronous thalamocortical activity (Llinas and Ribary, 1998). By recording cortical event-related potentials in a monkey during a visuomotor pattern discrimination task, Ding et al. (Ding et al., 2000) discriminated three different coordination states, each lasting around 100 ms with quick transitions between them. Laminar recordings in monkeys revealed transient beta bursts lasting about 150 ms (Lundqvist et al., 2016) related to memory encoding and decoding processes (Sherman et al., 2016). In human EEG and magnetoencephalographic (MEG) resting-state recordings, periods of oscillation bursts lasting around 200 ms have been described in the alpha (Williamson et al., 1996) and beta-bands (Seedat et al., 2020). Hidden Markov Models on MEG resting-state activity revealed short-lived transient brain states lasting around 50-100 ms, with spatially distinct power and phase-coupling in specific frequency bands (Vidaurre et al., 2018). Recently, using measures of entropy and hierarchy of functional magnetic resonance imaging (fMRI) signals, Deco et al. (Deco et al., 2019) demonstrated that the optimal timescale for discovering relevant spatiotemporal structures of brain signals is around 200 ms.

Overall, ample evidence indicates that spontaneous brain activity is parcelled into blocks of stable brain states that last around 100-200 ms, potentially representing the basic building blocks of consciousness. An increasingly popular method to capture these transient brain states are the EEG microstates, which have been suggested to represent the neural correlates of the elementary building blocks of the contents of consciousness, the “atoms of thought” (Baars, 2002a; Bressler and Kelso, 2001; Changeux and Michel, 2004; Lehmann et al., 1987). The concept of EEG microstates was developed three decades ago from the purely phenomenological observation that the head-surface voltage topographies recorded with multichannel EEG do not randomly change in space and time. Rather, a small set of prototypical topographies exist that remain quasi-stable for about 50-150 ms and rapidly transition from one to the other (Creaser et al., 2021; Lehmann et al., 1987). These transiently stable topographies emerge from the temporary synchronized neuronal activity of large-scale networks (Koenig et al., 2002; Michel and Koenig, 2018).

Several studies showed the stability of the dominant microstate topographies within and across subjects, independent of age and gender (Jabèsa et al., 2021; Koenig et al., 2002; Tomescu et al., 2018; Zanesco et al., 2020). However, the temporal dynamics of the microstates, such as frequency of occurrence, duration, or transition probabilities, are susceptible to the momentary state of the brain. For example, instructing participants to focus their thoughts on specific autobiographical memories, on previously seen objects, on the definition of particular words, or arithmetic calculations selectively influence duration or occurrence of specific microstates (Bréchet et al., 2019; Milz et al., 2016; Seitzman et al., 2017). Also, meditation leads to the alteration of the presence of distinct microstates (Brechet et al., 2021; Faber et al., 2017; Panda et al., 2016; Zanesco et al., 2021). Most importantly, different neuropsychiatric and neurological diseases, particularly schizophrenia, have been shown to alter the temporal characteristics of specific EEG microstates(Andreou et al., 2014; Kindler et al., 2011; Lehmann et al., 2005; Rieger et al., 2016; Strelets et al., 2003; Tomescu et al., 2015).

While these and many other studies demonstrated the sensitivity of EEG microstate dynamics to the momentary mental or cognitive state of the healthy and the pathological brain, little is known about the changes of EEG microstates due to altered states of consciousness such as sleep, anesthesia, or clinical conditions like coma or minimally conscious states. If EEG microstates are indeed the neural correlates of the elementary building blocks of the contents of consciousness, then any alteration of the consciousness level should modulate EEG microstates, either in terms of the diversity of states, the duration of a given state, or the syntax of transition between different microstates. The few existing studies indicate such modulations. Sleep as compared to wakefulness did not alter the topography of the most dominant microstates, but in a deep sleep (stage N3), the duration of all microstates increased (Brodbeck et al., 2012). A more recent study with high-density EEG source imaging demonstrated an increased presence of two EEG microstates during non-rapid eye movement (NREM) sleep compared to wakefulness, associated with low-frequency activity in the medial frontal and the occipital/thalamic regions, respectively (Bréchet et al., 2020). Another recent study (Gui et al., 2020) used the EEG microstate approach to assess residual cognitive functions in unresponsive patients. The authors showed a reduction of microstates that are thought to reflect higher cognitive functions, while microstates associated with basic sensory functions were increased compared to controls. They also showed that the duration and occurrence of the “cognitive” microstates reflected the strength of residual consciousness and predicted recovery in these patients. Similarly, the ability of microstate temporal parameters to predict recovery from the coma has been demonstrated in (Stefan et al., 2018).

To the best of our knowledge, only one study used the EEG microstate approach to study the effects of mild to moderate sedation induced by anesthetics(Shi et al., 2020), but did not examine the different loss of consciousness conditions. The authors showed that one specific microstate became salient during moderate sedation induced by propofol in increased time coverage and occurrence.

In this study, we investigated the spatio-temporal properties of EEG microstates by following surgical patients from the awake condition to loss of consciousness and further to surgical anesthesia. As an anesthetic, we used intravenous propofol, which is a widely used, short-acting GABA receptor agonist. To provide surgical anesthesia, we used supplemental sufentanil, which is a strong opioid, and rocuronium, which is a muscle relaxant. We aimed to highlight the difference between fully alert/baseline compared to surgical anesthesia conditions and the actual correlations of brain dynamics during that transition to advance further our understanding of conscious and unconscious states of the human brain. We also introduce a new method to evaluate the temporal sequence of EEG microstates, based on the algorithmic Lempel-Ziv complexity index which, contrary to what has been described previously (Casali et al., 2013; Schartner et al., 2015), is based on the temporal dynamics of the whole scalp potential field rather than binarized EEG single-channel envelopes. The measure is holistic as it involves all EEG channels. It is reference-free as it is based on the spatial configuration of the potential field, in contrast to single-channel waveform analysis (Michel and Murray, 2012). Other authors have computed the Lempel Ziv complexity (PCI) of TMS-induced EEG activation patterns to assess the level of consciousness (Casali et al., 2013; Casarotto et al., 2016; Comolatti et al., 2019). It has been proposed that this index quantifies the brain’s ability of information integration after stimulation (Tononi et al., 2016). The need for TMS stimulation to determine the evoked EEG complexity arises however from the lack of control of PCI when applied directly on resting state data as even a tiny fraction of noise would increase entropy, reducing the generalizability of the measure. By relying on the very well-established microstate extraction procedure, our EEG complexity measure can be applied to resting state data without the need for external stimulation and allows to further assess how the different levels of pharmacologically-induced, altered states of consciousness influence the complexity of the microstate temporal dynamics.

## Materials and Methods

### A. Experiment protocol

Twenty-three patients scheduled for minor elective surgery (ear-nose-throat/plastic surgery) were included at the time of the anesthesia consultation after giving written informed consent. The study protocol was approved by the Ethics Committee of Geneva University Hospitals (CER 12-280). None suffered from current or prior neurological or psychiatric impairments. The complete list of inclusion and exclusion criteria is available in the **Appendix**. The mean age of participants was 30 years (range 20-47 years). No monetary compensation was offered.

The patients fasted for at least six hours before anesthesia for solids and two hours for clear liquids (Smith et al., 2011). They did not receive any preoperative anxiolysis. On arrival in the operating theatre, standard non-invasive monitoring was installed, including a three-lead electrocardiogram (ECG), blood pressure cuff, end-tidal CO_2,_ and peripheral pulse oximetry. Oxygen (100%) was administrated through a facemask throughout the study period.

Propofol, prepared by the anesthesia team, was administered using a Target Controlled Infusion (TCI) device (Base Primea, Fresenius-Vial, Brezins, France) and the pharmacokinetic-pharmacodynamic (PK/PD) model by Schnider et al. (Schnider et al., 1999). The TCI device estimates the propofol concentrations in the plasma and at the effect-site (brain). The initially chosen effect-site concentration was 0.5 μg ml^-1^. We assumed that the equilibration of the blood-brain concentrations (“steady-state”) was achieved within 5 minutes after identical plasma and cerebral concentrations appeared on the TCI device screen. Effect-site concentrations were then increased stepwise by 1.0 μg ml^-1^ until 2.5 μg ml^-1^ and then by 0.5 μg ml^-1^ until loss of consciousness (LOC). After each increase, the “steady-state” was maintained for 5 minutes, and after this period, a five-minute EEG recording was done.

To estimate the degree of alertness, from fully alert to surgical anesthesia, we used a modified Observer’s Assessment of Alertness/Sedation (OAA/S) scale. The original OAA/S scale was developed to evaluate the depth of sedation clinically and to identify the time point of LOC in patients receiving sedative drugs (Chernik et al., 1990). The 5-point scale ranges from 5 (fully awake = subject responds readily to name spoken in a normal tone, normal speech, normal facial expression, no ptosis) to 1 (deep sedation = subject does not respond to mild prodding or shaking) (**Table 1**). OAA/S 5 was called BASE (baseline), and LOC was defined as a score ≤2. Increasing depths of sedation from OAA/S 5 to OAA/S 1 were achieved exclusively with increasing effect-site propofol concentrations and without any additional medication. However, as OAA/S 1 states (subject not responding to mild prodding or shaking) does not ensure that the subject does not react to active facemask ventilation (which may interfere with EEG recordings) and does not correspond to surgical anesthesia, we added a further state, called DEEP. DEEP was achieved in further increasing the depth of sedation at OASS 1 through adding supplementary propofol to eventually reach effect-site concentrations between 4 and 5.5 mcg/ml, and additionally, an intravenous bolus dose of each, a strong opioid (sufentanil 0.2 mcg/kg) and a neuro-muscular blocking agent (rocuronium 0.6 mg/kg). Rocuronium was administered to counteract a potential sufentanil-related muscle rigidity and to facilitate oro-tracheal intubation. EEG recordings were ended before oro-tracheal intubation.

**Table 1.**
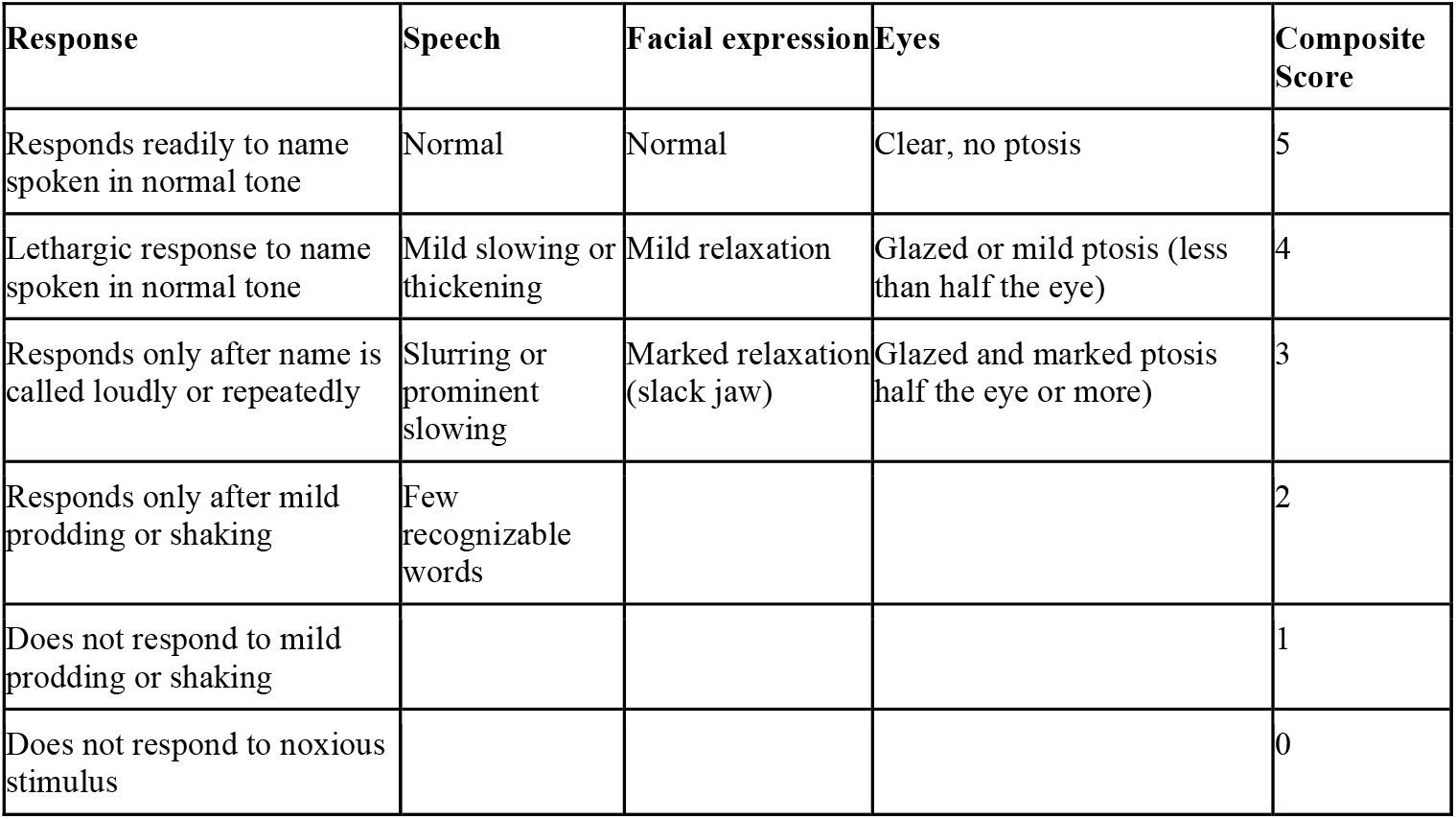
Observer Assessment of Alertness/Sedation (OAA/S) scale. Correspondence between Response, Speech, Facial Expression, Eyes characteristics and Composite score according to the Observer Assessment of Alertness/Sedation (OAA/S) scale

During the anesthesia procedure, vital signs were continuously recorded using the institutional computerized anesthesia record chart. They included heart rate, systolic, diastolic, mean arterial blood pressure, oxygen saturation (pulsoxymetry), and end-tidal CO_2_. These variables were not used for analyses. Neural correlates of propofol-induced anesthesia were investigated by acquiring 64-channel electroencephalographic (EEG) data with active Ag/AgCl electrodes (actiCap; BrainProducts) in an extended 10-10 System under the control of neuroscientists (Oostenveld and Praamstra, 2001). Prior to the arrival in the operation room, subjects were instructed to stay with closed eyes and to relax as much as possible. After a resting period of 10 minutes, a baseline EEG (BASE) was recorded (5 minutes duration). The 5 minutes EEG recording was repeated at each propofol state with a band pass filter between DC and 1000 Hz and was digitized at 5 kHz, with an online reference at FCz.

### B. The path to unconsciousness and surgical anesthesia

Each dataset was annotated every minute with a level of sedation going from BASE to DEEP. Since starting from BASE, every subject reached LOC and eventually DEEP, it was possible to define a “path from consciousness (fully alert) to surgical anesthesia” as the series of conditions traversed by the subject. **Fig. 1** represents such a path for the subjects. **Fig. 1, Panel A** shows, for each condition, the distribution of propofol effect site concentrations across the whole population and within each condition. Each box plot shows the average distribution across the population, the values corresponding to the 25^th^ and 75^th^ percentile of the distribution, and the maximum and minimum values that are not outliers. Conversely, **Fig. 1, Panel B** represents the Violin distribution of conditions across the whole population with respect to the measured propofol effect-site concentrations.

**Fig. 1.**
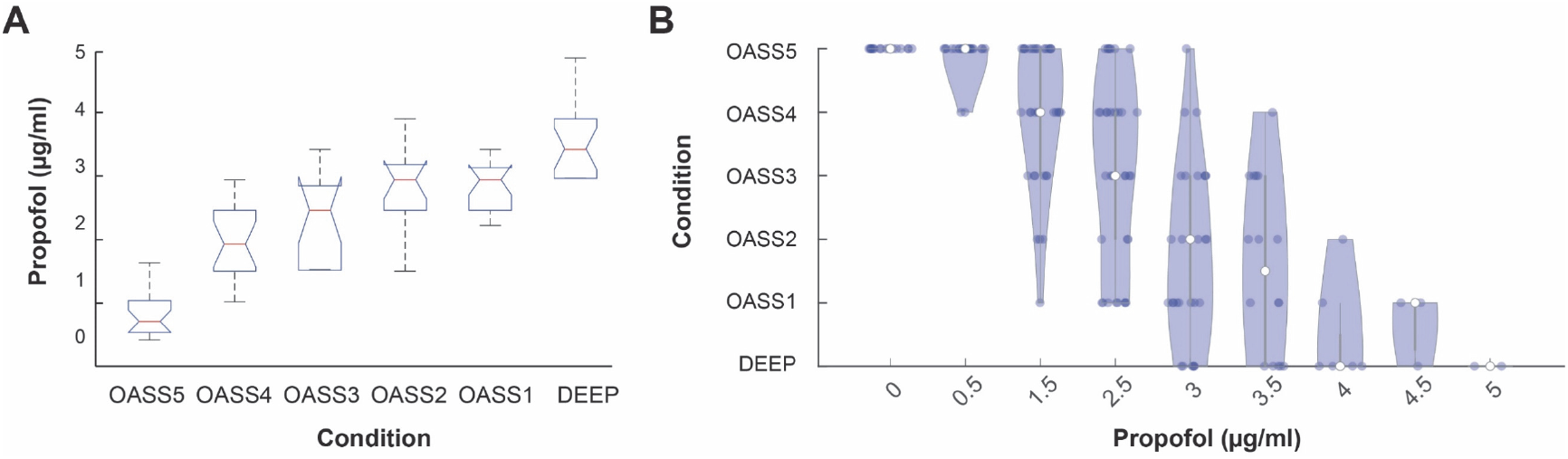
Behavioral data analysis. **Panel A**. Box-plot representation of the “*path to unconsciousness*” taken by each subject (each blue line) to reach condition “DEEP” from “OASS5” (x axis) in relation to Propofol concentration (y axis). Each patient was infused with a steadily increasing dose of Propofol with plasma concentrations ranging from 0.5 μg/ml to 4.5 μg/ml. Every minute while Propofol was infused, expert clinicians performed a clinical assessment of consciousness according to the Observer Assessment of Alertness/Sedation scale (OAA/S) with scores from 1 to 5 (indicated in the picture as “OASS1”, “OASS2”, …, “OASS5”). Conditions of deep anesthesia-induced loss of consciousness and baseline (subject fully awake, before any infusion of Propofol) are named “DEEP” and “BASE” respectively. **Panel B**. Violin plot distribution of the conditions of unconsciousness (from “OASS5” to “DEEP”) reached by each subject depending on the propofol level concentration (x axis, ranging from 0.5 to 4.5 μg/ml).

### C. The first stage of data preprocessing

Data were preprocessed with custom MATLAB scripts based on routines from the EEGLAB toolbox (Delorme and Makeig, 2004) and within Cartool (Brunet et al., 2011) in two stages (**Fig. 2**), following an increased-stability procedure also tested in previous works (Artoni et al., 2017). Within the first stage (**Fig. 2, STEP 1**), continuous data were processed using a Reliable Independent Component Analysis (RELICA) approach (Artoni et al., 2014) to remove artifacts and other non-neural noise sources, without any preliminary data dimensionality reduction (Artoni et al., 2018). To maximize both stability (Artoni et al., 2014) and dipolarity (Delorme et al., 2012) of Independent Components (ICs), raw data were first high-pass filtered using a zero-phase 1.2Hz, 24^th^ order Chebyshev type II filter, then low-pass filtered using a zero-phase 45Hz, 70^th^ order Chebyshev type II filter and resampled at 250Hz. Thanks to the rollover steepness of the filter, there was no need to perform a further 50Hz Comb Notch Filter. Channels having Kurtosis outside 5 standard deviations with respect to other channels or having prominent prolonged artifacts as confirmed by visual inspection were removed. Epochs with high-amplitude artifacts or high-frequency muscle noise were also identified by visual inspection and removed. The remaining data were submitted to RELICA with an AMICA core (Artoni et al., 2014) and 100 point-by-point Infomax ICA with a GPU-accelerated BeamICA implementation (Kothe and Makeig, 2013). Final ICA mixing and unmixing weights were then collected.

**Fig. 2.**
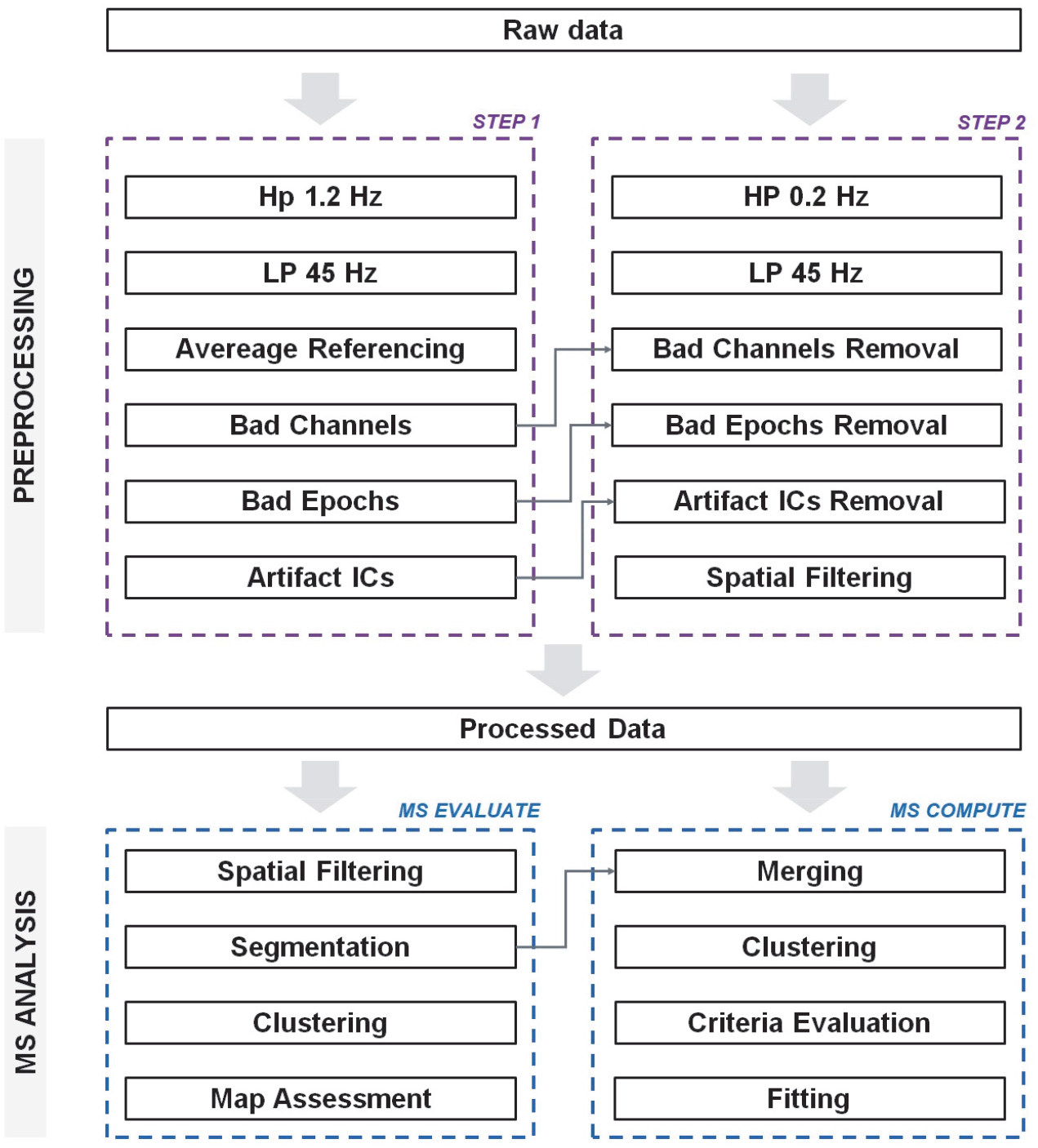
Preprocessing and Analysis pipeline. Schematics representing the steps performed for preprocessing and subsequent microstate analysis (see “Methods”). **STEP 1** includes relative aggressive filtering steps with the aim of determining, for each subject, bad channels, bad epochs, and artifact Independent Components (ICs) to remove. **STEP 2** includes more conservative filtering and bad channels. Bad epochs and artifact ICs are removed according to the information retrieved from STEP 1 before performing spatial filtering and entering the microstates (MS) analysis pipeline. In the MS EVALUATE pipeline, processed data from each subject goes through the classical steps of Spatial Filtering and Segmentation, pooled data for all subjects and each condition individually are clustered and maps assessed for similarity across conditions. Next, in the MS COMPUTE pipeline, pooled segmentation data for all subjects and conditions are merged, clustered and evaluated according to multiple state of the art criteria (see “Methods” and **Fig. 7**). Final MS maps are then fitted to each single-subject dataset to obtain various features such as coverage, global explained variance, MS-wise power etc.

### D. The second stage of data preprocessing

Within the second preprocessing stage (**Fig. 2, STEP 2**), raw data were high-passed using a zero-phase 0.2Hz 24^th^ order Chebyshev type II filter and a zero-phase 45Hz, 24^th^ order Chebyshev type II Low Pass filter and resampled at 250Hz. Bad channels and epochs already identified within the first preprocessing stage were rejected, and data were carefully visually re-inspected for any remaining artifacts. ICA unmixing weights computed within the first preprocessing stage were then re-applied to the dataset, and source localization was performed using the Dipfit toolbox (Delorme et al., 2012) within EEGLAB. Dipolar and stable ICs related to stereotyped artifacts such as eye activity and neck muscle activity were removed from the data by back projecting the IC activation data after zeroing out the columns of the mixing matrix corresponding to the artifact ICs. Missing channels were interpolated, and clean data were spatially filtered within Cartool to improve the SNR of the data (Michel and Brunet, 2019).

### E. Extraction of microstates

EEG microstate segmentation was performed using the standard procedure also described in (Murray et al., 2008), while taking extra precautions to ensure the statistical reliability of all the results. In fact, throughout all the analyses, excluded time epochs (beginnings and ends) were treated as “boundaries”, that is “gaps” in the data that could not be “crossed” by the analysis steps. First, the Global Field Power (GFP) maxima were extracted from each participant’s spontaneous preprocessed EEG. For each condition and participant, the GFP peak maps (channel values at the timestamp corresponding to the GFP peak) were extracted to ensure a high signal-to-noise ratio (Koenig et al., 2002) and were clustered via modified k-means to extract distinct templates (Murray et al., 2008; Pascual-Marqui et al., 1995). Within this step, the spatial correlation between each GFP map and each template randomly generated was calculated while ignoring the polarity of maps (Michel and Koenig, 2018).

Each template was iteratively updated by averaging the GFP maps that presented the highest correlation with the template. At the same time, the Global Explained Variance (GEV) of template maps was calculated, and the process was iterated until the stability of GEV was reached. For each condition (BASE through DEEP), the optimal number of microstate classes was determined using different criteria (**Fig. 3**), each estimating the “quality” of a single segmentation according to specific metric, namely “Gamma”, “Silhouettes”,” Davies-Bouldin”, “Point-Biserial”, “Dunn-Robust” and “Krzanowski-Lai” (KL), the optimum subsequently validated by a MetaCriterion implemented and published with the Cartool toolbox and discussed in depth in (Bréchet et al., 2019).

**Fig. 3.**
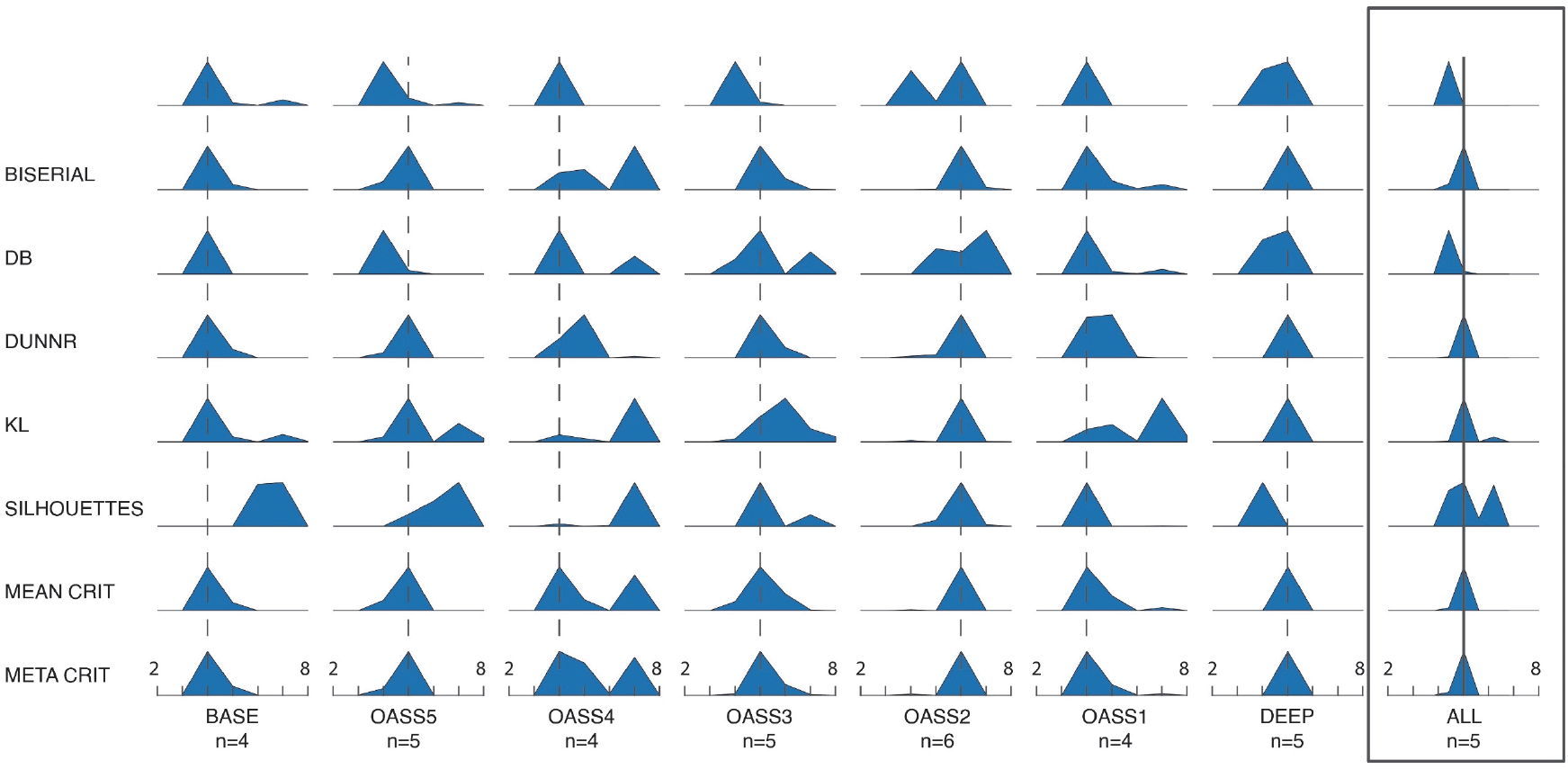
Microstates assessment criteria. Assessment of extracted microstates with different criteria, each estimating the “quality” of a single segmentation, according to specific metrics. The criteria used were “Gamma” (GAMMA), Point-Biserial (BISERIAL), Davies-Bouldin (DB), Dunn Robust (DUNNR), “Krzanowski - Lai” (KL), “Silhouettes (SILHOUETTES)”. These criteria were also combined into a “Mean criterion” (MEAN CRIT), that is a criterion representing the “average” of the probabilities yielded by all the other criteria and the “Meta criterion” (META CRIT) that represents the best principled choice for the number of microstates. For each condition analyzed (columns, “BASE” through “DEEP”) and assessment criteria (“GAMMA” through “META CRIT”) a plot shows the probability (according to the specific criterion) for each number of microstates (2 to 8). A dashed vertical line for each column represents the best number of microstates according to the metacriterion for the relative condition. The rightmost column represents the criteria probability for the pooled dataset (“ALL” conditions), with the metacriterion unequivocally suggesting n=5 microstates.

The dominant microstates were identified within each condition from the templates across participants using a second modified k-means clustering step. Each clustering step was computed 100 times to maximize stability and to overcome the possible statistical instability of the randomization procedure within the k-means algorithm (Murray et al., 2008). The spatial correlation was finally computed between microstates across all conditions. Each microstate was labeled as the name of the microstate in baseline with minimal topographical dissimilarity (i.e., highest spatial correlation).

### F. Assessment of the quality of microstates and fitting

The topographical dissimilarity across different microstates both within each condition and across different conditions was also computed to ensure no “microstate splitting” occurred. **Fig. 4, Panel A** reports the topographical correlation values between DEEP and BASE microstates. Average and standard deviations of the spatial correlation of paired microstate maps across all conditions are finally reported at the bottom of **Fig. 4, Panel B**. Given the high correlation between paired maps across conditions and the similar assessment of the optimal number of microstates yielded by the meta-criterion, the data of all conditions were pooled (condition ALL) and the hitherto described analysis repeated as shown in **Fig. 2**.

**Fig. 4.**
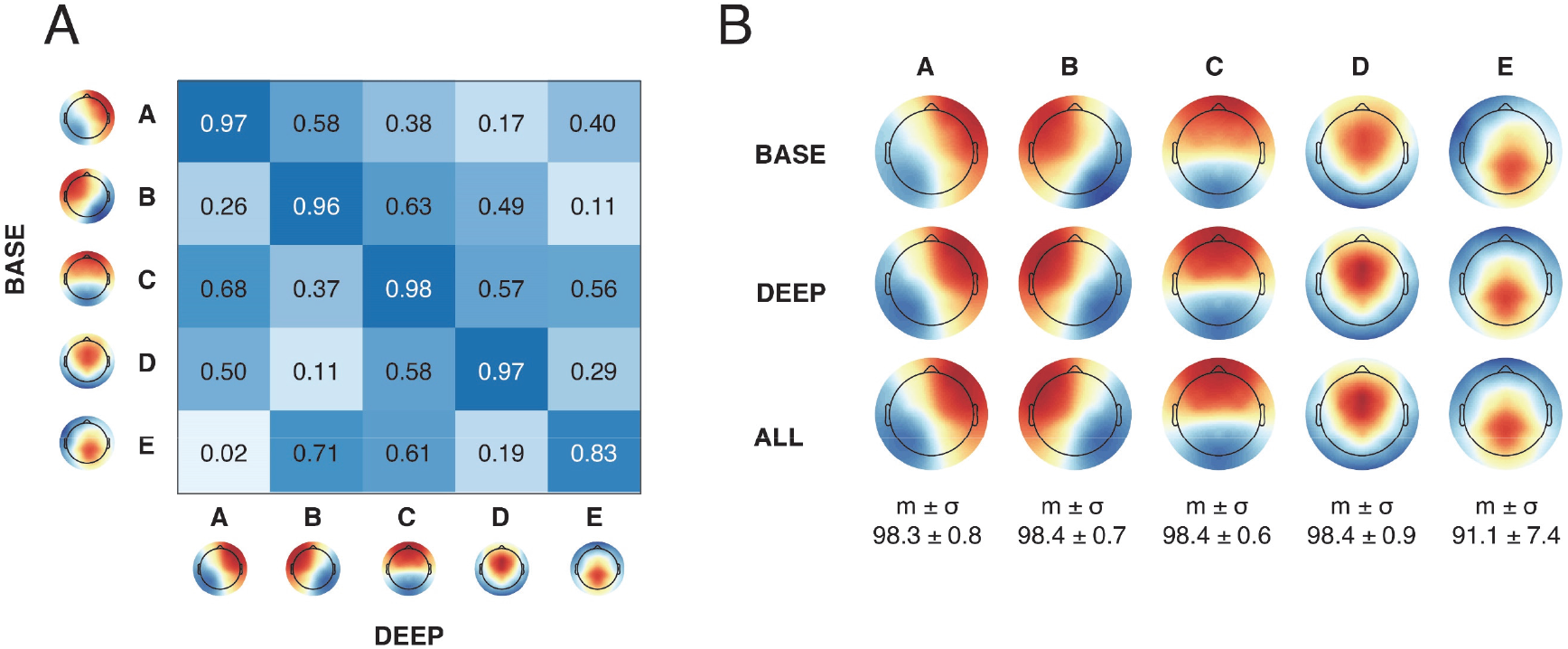
Microstates assessment. **Panel A**. Similarity matrix (absolute correlation coefficient) representing the absolute correlation (rounded to the 2^nd^ digit) of microstates maps between BASE and DEEP conditions. **Panel B**. Representation of the 5 microstates obtained respectively for conditions “BASE”, “DEEP” and “ALL”. The average correlation of ordered microstates maps (all possible pairs) and the standard deviation are reported at the bottom of each column (microstates from A to E).

Finally, spatial correlation between the templates identified at the group level (ALL) and those identified for each subject was computed using a temporal constraint (Segments Temporal Smoothing) of 6 samples (24 ms). EEG frames were labeled in a “winner-takes-all” strategy (Michel and Koenig, 2018) according to the group template it best corresponded to (no labeling was performed at correlations lower than 0.5), which generated the microstate sequence for further analysis.

### G. Extraction of microstates canonical features

For each condition and each subject, the following microstate features were computed:

- Global explained variance (GEV), obtained for each microstate class, as the sum of the explained variances weighted by the Global Field Power at each time point, that is

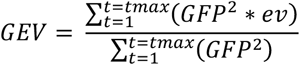
- Spatial correlation, obtained for each microstate class, as the mean spatial correlation of the microstate map with the GFP peak maps within the spatially filtered dataset. The spatial correlation between two maps is mathematically defined as 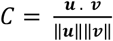 where ***u*** and ***v*** represent the first and second map respectively and ||.|| the *𝓁*2 norm.
- Duration, obtained for each microstate by averaging the time said microstate is active (in a winner-takes-all fashion) before transitioning to another microstate.
- Density, computed for each microstate as the number of occurrences of said microstate per second of data.
- Coverage, computed for each microstate as the relative number of time points of the dataset covered by said microstate

### H. Statistical comparison of microstate features across conditions

After rejecting the null hypothesis of data Normal distribution over each group using a Kolmogorov Smirnov test (significance set at *α* = 0.05), these measures were compared across conditions using a Kruskal Wallis test followed by a Tukey’s Honest Significant Difference (HSD) criterion for post-hoc comparison. In the following, median (MED) and 95% confidence interval of the median (STM) are reported instead of the mean (AVG) and standard deviation (STD) whenever data did not follow a standard distribution. STE is reported as 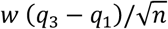 with *w* = 1.57, *q*_3_ and *q*_1_ the 75^th^ and 25^th^ percentile, respectively, and *n* the number of samples (Chambers et al., 1983). Violin plot distributions were also calculated by kernel density estimation with a Gaussian kernel to minimize the *𝓁*2 mean integrated squared error (Silverman, 1986). Box plots with comparisons across all conditions for the most significant measures (Density, Duration, Spatial Correlation) are represented in **Fig. 5**.

**Fig. 5.**
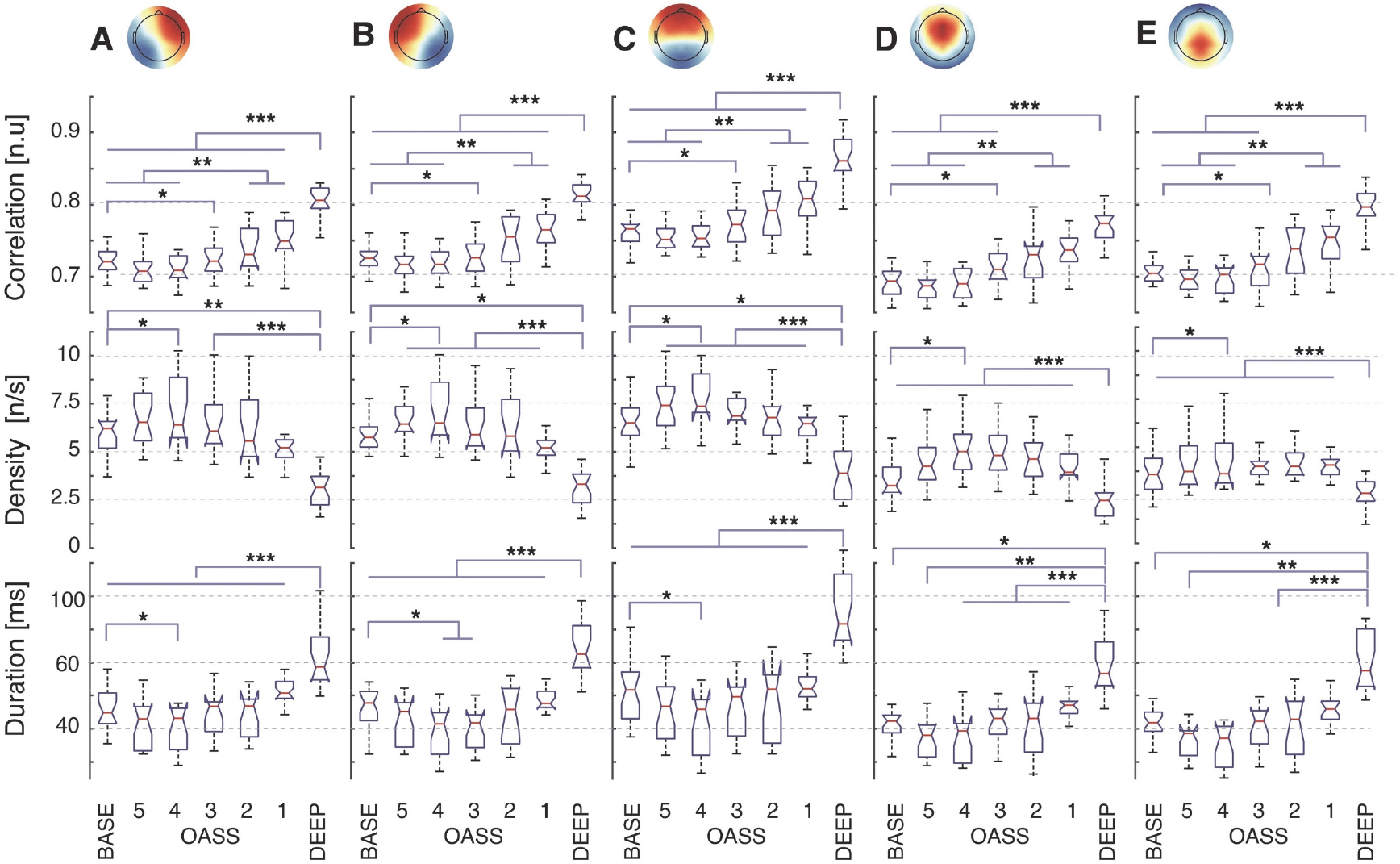
Detailed microstate features comparison across conditions. Detailed comparison boxplot of the most important features (from top to bottom: Correlation, Density, Duration) across all conditions (BASE through DEEP) and for each microstate (A through E). On each box, the central red mark indicates the median of the distribution, the bottom and top edges of the box indicate the 25^th^ and 75^th^ percentiles, respectively. The whiskers extend to the extreme data points (roughly corresponding to 99.3% of data if they are normally distributed) that are not considered outliers. Points within the distribution are considered outliers if greater than *q*_3_ + *w*(*q*_3_ - *q*_1_) where *q*_1_ and *q*_3_ are the 25^th^ and 75^th^ percentiles of the sample data, respectively and *w* the maximum whisker length. * p<0.05; ** p<0.01; *** p<0.001.

### I. Checking for polarity inversions to test for nonlinearity

After preliminary observations of the data, the possibility of a nonlinear path to unconsciousness was tested by computing for each microstate the relative normalized percentage difference of significant features (Density, Duration, Spatial Correlation) in OASS4 and DEEP conditions with respect to BASE (**Fig. 6**), each relative difference was statistically tested against a null distribution. An inversion of polarity between the first and second bar groups demonstrates a nonlinear path to unconsciousness.

**Fig. 6.**
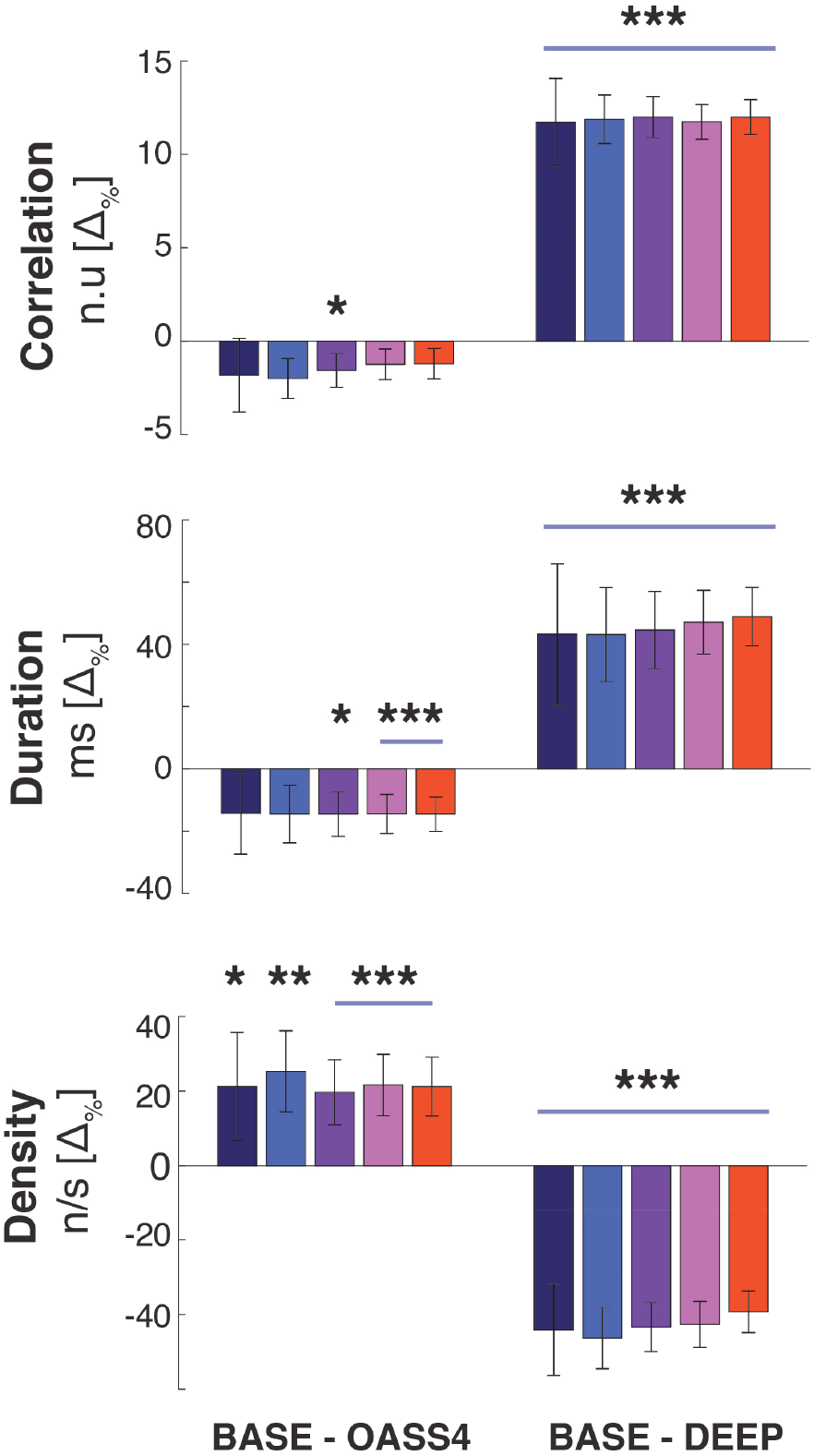
Nonlinearity of the path to unconsciousness. Relative percentage difference of features (Correlation, Density, Duration) between conditions OASS4 and BASE (leftmost bar group) and between conditions DEEP and BASE (rightmost bar group), calculated for each microstate (A through E, color coded with colors ranging from Blue to Red). A significant inversion of polarity between the first and second bar groups demonstrates a significantly nonlinear path to unconsciousness: Correlation and Duration decrease with respect to BASE before reaching a maximum in DEEP while Density increases with respect to BASE before reaching a minimum in DEEP.

### K. Complexity analysis

To compute the complexity of non-binarized sequences, we used the Lempel-Ziv-Markov chain algorithm (LZMA2) for lossless data compression (Pavlov, 2013a) with maximum compression level, 64MB dictionary, 64 FastBytes, BT4 MatchFinder, BCJ2 Filter (Pavlov, 2013b). The Microstate Lempel-Ziv Complexity (MS-LZC) is defined as the compressed size (in byte) of a microstate sequence. For each subject and condition, the MS-LZC was computed for each extracted window from the full microstate sequence, using a sliding-window approach (5s window length, 4s window overlap) to ensure a smooth and representative output. The MS-LZC for windows overlapping with two or more conditions were discarded to avoid discontinuities. MS-LZC for each subject was divided by the baseline MS-LZC amplitude (BASE condition). The grand average MS-LZC for each condition was obtained by averaging the normalized MS-LZC across subjects (**Fig. 7, Panel A**). Statistical comparisons across conditions were then performed here in a similar way as explained in **Section “L”** above. A different representation of MS-LZC over time was obtained instead by averaging the LZC time course across subjects after time-warping each to a common median length (**Fig. 7, Panel B**)

**Fig. 4.**
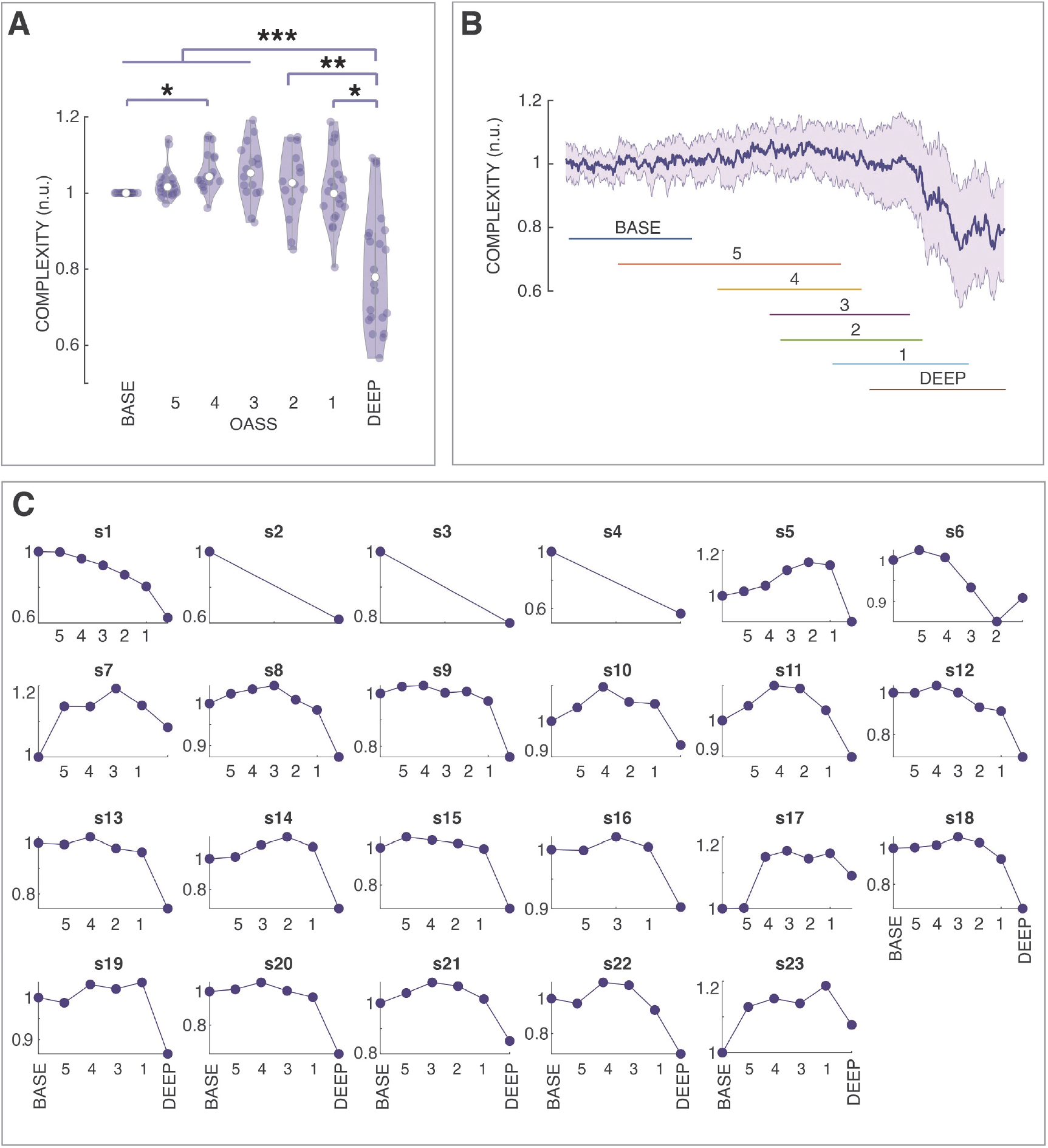
Lempel-Ziv Complexity Analysis of Microstate sequences. **Panel A**: violin distribution of the normalized Microstate Lempel-Ziv Complexity (MS-LZC) of microstates across subjects for each OASS condition (BASE through DEEP) and statistical significance of comparisons (* p<0.05; ** p<0.01; *** p<0.001). **Panel B**: Grand-average (bold blue line) and standard deviation (blue shading) of the time-warped MS-LZC across subjects. The colored lines represent the minimum and maximum span in the common warped time axis for each condition (BASE – OASS5,4,3,2,1 - DEEP). **Panel C**: Single-subject average normalized MS-LZC for each condition

## Results

### A. From fully alert to surgical anesthesia

Twenty-three patients scheduled for minor elective surgery volunteered for the experiment. Their consciousness state was assessed with the Observer Assessment of Alertness/Sedation (OAA/S) from fully awake with initiated Propofol injection, to OASS0/DEEP – fully anesthetized – **(Table 1**). All patients reached DEEP (surgical anesthesia) within 20 ± 6 minutes when infused with increasing target effect-site concentrations of propofol ranging from 0.5 μg/ml to 5.0 μg/ml, and additional sufentanil (and rocuronium) a soon as OASS 1 was reached (**Fig. 1**).

### B. Several criteria reveal five salient canonical microstates

Careful preprocessing (see **Methods**) and microstate analysis of the EEG data continuously collected during the experiment revealed five canonical microstates (named with letters A through E), best explaining the data according to a Meta-criterion (Brechet et al., 2020). Microstates (MS) had different spatial distributions. The spatial correlation across different microstates within each condition (e.g., MS A, and MS B within “BASE”) and across conditions (e.g., MS A for “BASE” and MS B for “DEEP) was always lower than the spatial correlation across paired microstates conditions (e.g., MS A for BASE and MS A for “DEEP”) (**Fig. 4, Panel A**). The average spatial correlation (x 100) across conditions of paired microstates was 98.3 ± 0.8 (microstate A), 98.4 ± 0.7 (microstate B), 98.3 ± 0.6 (microstate C), 98.4 ± 0.9 (microstate D), 91.1 ± 7.4 (microstate E) (**Fig. 4, Panel B**).

### C. Correlation, density, and duration successfully explain data

The maximum statistical separation between BASE and DEEP conditions was found for three microstate features (Correlation, Density, Duration). These features were compared across all OASS conditions, aiming not just to describe the starting and ending point but the whole path to unconsciousness (**Fig. 5**).

All features exhibit a nonlinear path to unconsciousness. The *spatial correlation* of the microstate maps fitted to the ongoing EEG in DEEP condition was significantly higher not only with respect to BASE but also regarding all other conditions – OASS 5, OASS 4, OASS 3, OASS 2, OASS 1 for microstates A, B and C (p<0.001); BASE, OASS 5, OASS 4, OASS 3 for microstates D and E. A significant difference was found between OASS 2, OASS 1, and BASE for all microstates (p<0.01). Interestingly, significant differences (p<0.05) could already be seen between BASE and OASS 3. However, although without reaching significance, the spatial correlation first decreased at the beginning of the path to unconsciousness, reaching the minimum at OASS 4 and then steadily increased until the maximum was reached with deep anesthesia.

Similarly, the *density* (number of occurrences of a microstate per second) of all microstates was lower for DEEP with respect to any other condition. However, as for the other microstate features, microstate density first increased at the beginning of Propofol administration before it decreased. This initial increase reached significance for all microstates at OASS 4 when compared to BASE.

The *duration* of all microstates was significantly higher for DEEP with respect to all other conditions (BASE through OASS 1), with p<0.001 for microstates A, B, and C. As for the other parameters, a U-shape behavior was observed with an initial decrease of the duration up to OASS 4, reaching significance (p<0.05) for microstates A and B compared to BASE, before steadily increasing from OASS 4 through DEEP.

### D. A significantly nonlinear path from fully alert to surgical anesthesia

All three microstate features showed a U-shaped behavior from awake to deep anesthesia. To further explore this phenomenon, relative normalized differences of the microstate features were calculated (**Fig. 6)**. This difference was negative for BASE – OASS 4, significant for the *spatial correlation* (Microstate C: p<0.05) and *duration* (microstate C: p<0.05, microstates D and E: p<0.001), and positive for BASE – DEEP, significant (p<0.001) for *spatial correlation* and *duration* for all microstates. On the contrary, microstate *density* exhibited a positive BASE – OASS 4 difference (microstate A: p<0.05, microstate B: p<0.01, microstates C,D,E: p<0.001) and negative BASE – DEEP difference (all microstates p<0.001), suggesting a nonlinear path to unconsciousness from BASE to DEEP (first a decrease, then an increase for spatial correlation and duration, vice versa for density).

### E. “Mild” and “deep” sedation respectively increase and decrease microstate complexity

Finally, based on the microstates sequence, we calculated a novel feature, the Microstate Sequence Lempel Ziv Complexity (MS-LZC), that captures the level of compressibility (size reduction) of a fixed-length sequence, expressed in kbit/s. An increase in this complexity value would indicate a more heterogeneous succession of the different microstates, while decreased complexity would reflect simpler and repetitive microstate sequences. Grand-average MS-LZC analysis (**Fig. 7, Panel A**) revealed a nonlinear path to unconsciousness with a MS-LZC increase (from BASE to OASS 4 and OASS 3) followed by an MS-LZC decrease (OASS 3,2,1, DEEP). Significance (see methods) was reached (i) between BASE and OASS 4 (p<0.05), (ii) between DEEP and BASE, OASS 5,4,3 (p<0.001), (iii) between DEEP and OASS 2 (p<0.01), (iv) between DEEP and OASS 1 (p<0.001). Complexity over time (**Fig. 7, Panel B**) also confirms a sustained MS-LZC, decreasing steadily towards DEEP. A nonlinear behavior was observed for all subjects (except s2, s3, s4, where conditions were not annotated), with a MS-LZC increase followed by a MS-LZC decrease. However, three of them (s7, s17, s23) reached a higher complexity during DEEP condition compared to BASE.

## Discussion

By applying the new method of microstate sequence complexity (MS-LZC), along with the classical microstate features, our study revealed a distinct “U-shaped” path of propofol-anesthetized patients from fully alert (baseline) to surgical anesthesia. Our results demonstrate the value of microstates in capturing and synthesizing complex dynamical features of whole-brain networks in the sub-second time range that characterizes different states of consciousness.

Interestingly, we found a reversal effect of propofol from baseline to light sedation and from sedation to surgical anesthesia. This peculiar behavior is probably linked to the paradoxical excitation effect of propofol and other anesthetics at a lower dose (Ching et al., 2010; McCarthy et al., 2008), marked by disinhibition, loss of affective control (Fulton and Mullen, 2000), and seizure-like phenomena ranging from involuntary movements to generalized tonic-clonic seizures (Walder et al., 2002). In our study, this enhanced level of excitability is characterized by more diverse spatiotemporal EEG patterns (highlighted by shorter duration, higher density, and lower correlation of the microstates, as well as increased complexity of the microstate sequences). Therefore, these results suggest the presence of an intermediate brain state as compared to a fully awake condition, during which patients enter a state of hyperexcitability with increased complexity and probably greater awareness of both inside and outside stimuli. This state is also described under psilocybin as the “entropic brain” state (Carhart-Harris et al., 2014). Interestingly, this hyperexcitability state seemed to protract until full loss of consciousness in three subjects (**Fig. 7, Panel C**).

By further increasing propofol dosage, the level of excitability decreases and preludes complete loss of consciousness. In terms of EEG microstates, this effect is characterized by increasing duration, increasing correlation, decreasing the occurrence of the microstates, and reducing the complexity of the microstate sequences. These effects are reminiscent of the decreased excitability of the cortical networks due to enhanced GABAergic phasic and tonic currents induced by propofol (Dasilva et al., 2020; Orser et al., 1994). The effect was most pronounced at the stage of surgical anesthesia, where opioid analgesic sufentanil and the muscle relaxant rocuronium were added to reach complete unconsciousness.

The concept of fragmentation of consciousness in transiently stable brain states postulates that the non-continuous dynamics of these states are governed by “metastability” that hold the system in a critical balance and allow a flexible switch from one state to another (Deco and Jirsa, 2012; Jirsa et al., 1998; Tognoli and Kelso, 2014; Tononi et al., 1994). Any disturbance of this critical temporal balance of brain states, being increased or decreased, might cause alterations in the global state of consciousness. Our finding of a U-shaped behavior of the temporal characteristics of microstates further underlines the importance of optimal metastability between order and chaos. Increased complexity of the network dynamics, as seen in OASS 4 and decreased complexity as seen in deep anesthesia, lead first to altered states of consciousness represented by a hyper-excitation and second to complete unconsciousness. Interestingly, in the case of induction of these states by propofol, all microstates were similarly affected. This effect is in contrast to sleep that selectively changes the temporal characteristics of specific microstates. A recent study (Bréchet et al., 2020) showed that two EEG microstates (a frontal and occipital/thalamic one) were highly represented during NREM sleep than resting wake state. However, dreaming during NREM sleep was associated with a decrease in the occipital/thalamic microstate presence, while the frontal microstate increased during dreams. The authors venture that reducing the occipital microstate slow-wave activity may indicate local activations that account for remembered dreams with rich perceptual content. In contrast, the increase of the frontal microstate may account for the executive disconnection of the dreaming brain to maintain sleep. Notably, these dreams were remembered and could be recalled means that memory is not entirely lost during sleep. This result is in contrast to propofol-induced anesthesia, during which memory is lost. It has been shown that the effect of propofol on memory is different from the sedative effect (Veselis et al., 2001). It might be that this amnestic effect explains why all microstates, and thus all functional brain networks, were similarly affected by propofol.

Surgical anesthesia results in a highly significant increase in the spatial correlation of the microstate template maps with the ongoing EEG (from 0.7 to 0.8 – **Fig. 5**). This result further highlights a decrease in complexity brought forward by deep anesthesia. Few spatial filters (i.e., 5 microstates) can better explain the ongoing EEG, in line with results suggesting a reduction in complexity to be a predictor of unconsciousness (Dasilva et al., 2020; Zhang et al., 2001). Considering that a winner-takes-all strategy was used to estimate microstate topographies (see **Methods**), an increased duration combined with increased spatial correlation is suggestive of longer-sustained and well-defined states slower-changing, simpler topographies, better correlated with smooth canonical microstate topographies. This effect could be attributed to the incidence of slow waves during propofol-induced anesthesia comparable to slow waves during sleep (Murphy et al., 2011). These low-frequency oscillations are associated with neuronal bi-stability and impaired network interactions caused by disruption of communication in and/or among cortical brain regions (Bellesi et al., 2014). It has been suggested that the presence of the microstates during low-frequency activity (such as during NREM sleep) reflects a temporary process of suppression of functional integration between the nodes of the network that generated the corresponding microstate, thus a deactivation of the network (Brechet et al., 2020). Following this interpretation, the increased duration of all microstates during a loss of consciousness reflects a continuous deactivation of the networks. In contrast to NREM sleep, all microstates were similarly prolongated and better presented, while in NREM sleep, only 2 microstates changed compared to wakefulness (Brechet et al., 2020). Only during deep sleep stage N3, the duration of all microstates increases (Brodbeck et al., 2012). This effect indicates that loss of consciousness during anesthesia is not similar to sleep.

Regarding the MS-LZC method proposed, previous studies have used the Lempel-Ziv compression (LZC) algorithm to evaluate the complexity and diversity of EEG signals, either with a single-channel or a multichannel approach (Casali et al., 2013; Schartner et al., 2015). In the single-channel approach, a raw or preprocessed/filtered EEG channel is divided into epochs, de-trended, and transformed with a Hilbert transform to estimate its envelope. The resulting signal is then binarized using a set threshold calculated as the mean value of the envelope itself: values of “1” and “0” are assigned to the time points respectively above or below said threshold, and the binarized sequence is then segmented into “binary words” by the LZC algorithm. The greater the number of “binary words”, with respect to the number obtained after randomly shuffling the original binarized sequence, the greater the complexity of the epoch. The multichannel approach is similar, with the only difference that binarized sequences obtained from each EEG channel are concatenated before submitting them to the LZC algorithm. This method (or others following a similar procedure), however, presents several drawbacks. First, both single-channel and multichannel approaches are highly influenced by the preprocessing of EEG data. In fact, noisy channels originate maximally-random sequences that can greatly reduce the robustness of the complexity measure. Second, envelope binarization is not representative of the data structure as both oscillations above and below the threshold are lost. Third, the method may result in a different binarizing threshold for each epoch, potentially leading to very different complexity values, even for contiguous epochs, if, for instance a particular event or burst of noise modifies the threshold to either very high or low levels. Fourth, with the multichannel approach, concatenation of binarized sequences from different channels may introduce discontinuities that may artificially increase estimated complexity. For example if sequence “A (0,0,0,0)”, with only one binary word of size 1, (“0”) is concatenated to sequence B (1,1,1,1), with only one binary word of size 1 (“1”), its concatenation A+B (0,0,0,0,1,1,1,1) would be composed by two binary words of size “1” and one artifact binary world of size 2 (“01”). Finally, and most importantly, single-channel analysis is highly dependent on the recording reference, while the topographic analysis used for microstate segmentation is completely reference-free(Michel and Murray, 2012).

This study has two main limitations. First, in the absence of an objective, clinical tool that allows quantification of the degree of unconsciousness, we were using the OAAS scale as a surrogate measure of alertness. The OAAS scale has been validated for sedative drugs, but it cannot be used to clinically measure the depth of surgical anesthesia. We, therefore, added an artificial score (DEEP) to the scale in deepening the degree of sedation to a status that empirically corresponded to surgical anesthesia. Based on daily clinical practice, we assumed that thanks to the combination of propofol and a strong opioid, surgical anesthesia was reached, although that state could not be quantified clinically. This also implies that, strictly speaking, our observations apply to different degrees of unconsciousness induced by propofol, with or without sufentanil. Second, the pathway from fully alert (BASE) to loss of consciousness (LOC) and then further to surgical anesthesia (DEEP) is a continuum. For instance, the end of “deep sedation” and the beginning of “surgical anesthesia” is not clearly defined. Also, this continuum is likely to depend on individual factors, such as a subject’s co-morbidities and the drugs used for sedation.

Overall, microstate sequence complexity and microstate features offer a granular and synthetic description that opens new perspectives on the neural correlates of transitions to loss of consciousness. The performance of current loss of consciousness decoders may be improved by considering the existence of a paradoxical excitation brain state, for example, by tracking the change of the slope in complexity rather than simply comparing features with the baseline condition. In future works, the microstate features and sequence complexity (see **Fig. 7, Panel C**) may also be used to track the path of recovery from loss of consciousness, explore possible relations to intra-operative awareness and sensitivity to propofol, or prevent propofol overdosing.

## Appendix

### Full inclusion/exclusion criteria

Subjects satisfied all of the following criteria to be enrolled in the study:

- Adult patients (age between 18 and 40 years)
- Right-handed
- American Society of Anesthesiology (ASA) status I-II
- Scheduled for elective surgery requiring a general anesthetic
- Able to read and understand the information sheet and to sign and date the consent form.

Subjects who potentially met any of the following criteria were excluded from participating in the study:

- Patients with significant cardio-respiratory or another end-organ disease (renal or hepatic disease influencing metabolism or elimination of study drugs).
- Patients with depression, neurological or psychiatry disorders.
- Dementia or inability to understand the informed consent.
- Patients with a history of esophageal reflux, hiatus hernia, or any other condition requiring rapid sequence induction of anesthesia.
- History of drugs (opioids) or alcohol abuse.
- Patients with a body mass index >30 kg m^-2^.
- Left-handed patients
- History of allergy or hypersensitivity to Propofol.

## Acknowledgments

The work was supported by the Swiss National Science Foundation (grant No. 320030_184677) and by the National Center of Competence in Research (NCCR) ‘SYNAPSY - The Synaptic Bases of Mental Diseases’ (SNF, grant n° 51AU40_125759). It was also supported by the Development Fund of the Medical Directories of the Geneva University Hospitals. We want to thank the research nurses, Ms. Béatrice Gil-Wey, Ms. Claudine Carera, and Mr. Patrick Huwiler, for their outstanding work. We acknowledge access to the facilities and expertise of the CIBM Center for Biomedical Imaging, a Swiss research center of excellence founded and supported by Lausanne University Hospital (CHUV), University of Lausanne (UNIL), Ecole polytechnique fédérale de Lausanne (EPFL), University of Geneva (UNIGE) and Geneva University Hospitals (HUG).

## Author Contributions

JM, CL, and MRT were responsible for patient recruitment and the clinical care of the patients during the recordings. JM, JB, CL, MRT and CMM designed the study. JB recorded the data. FA, developed the processing algorithms and analyzed the data. CMM supervised the analysis activities. MS and CMM contributed critical feedback to the data analysis and presentation of the results. FA, LB and CMM wrote the manuscript. All authors corrected and commented on the manuscript.

## Competing Interets

There are no competing interests pertaining to this manuscript

